## Supporting information for "Targeted chromosomal *Escherichia coli:dnaB* exterior surface residues regulate DNA helicase behavior to maintain genomic stability and organismal fitness"

**Table 1: Oligonucleotides**

| Name | Sequence (5’-3’) |
|---|---|
| DNA165 Cy3 | 5’ Cy3-TCCCACCCAACCCGACCGGCATCTAGTCTGGTAGCGTGAGCGAACGGACC |
| DNA180 | 5’ TGACGTCGCACACCGTGCTC |
| DNA181 | 5’ TTTTTTTTTTTTTTTTTTTTTTTTTTTTTTGAGCACGGTGTGCGACGTCAGCCGGTCGGGTTGGGTGGGA-BHQ |
| DNA180 Cy5 | 5’ TGACGTCGCACACCGTGCTC-Cy5 |
| CRISPR R74A FW | 5’ AAACGTCATATCTTTACTGAAATGGCGCGTTTGCG |
| CRISPR R74A RV | 5’ AAAACGCAAACGCGCCATTTCAGTAAAGATATGAC |
| CRISPR R164A FW | 5’ AAACTCGCACGACTTTCGGCAATTTTAAAGACGCG |
| CRISPR R164A RV | 5’ AAAACGCGTCTTTAAAATTGCCGAAAGTCGTGCGA |
| CRISPR K180A FW | 5’ AAACATTGCCGAAAGTCGTGCGAACAAAGACGAAG |
| CRISPR K180A RV | 5’ AAAACTTCGTCTTTGTTCGCACGACTTTCGGCAAT |
| CRISPR R328/9A FW | 5’ AAACCCGCCGTGTTCACGGGCAATACGGCGTGCGG |
| CRISPR R328/9A RV | 5’ 5AAAACCGCACGCCGTATTGCCCGTGAACACGGCGG |
| R74A HR oligo | 5’ CACACCGTCATATCTTTACTGAAATGGCAGCGCTGCAAGAAAGCGGTAGCCCTATCGATC |
| R164A HR oligo | 5’ GCGAAGATCTGCTGGATCTGGCTGAATCTGCAGTCTTTAAAATTGCCGAAAGTCGTGCGA |
| K180A HR oligo | 5’ TGCCGAAAGTCGTGCGAACAAAGACGAGGGCCCGGCGAACATCGCCGATGTGCTCGACGC |
| R328/9A HR oligo | 5’ CGCCAACGGAAGTGCGTTCCCGCGCGGCCGCTATTGCCCGTGAACACGGCGGCATCGG |

Base changes are underlined; Novel RE sights are highlighted in grey

| Table 2: Strains | | |
|---|---|---|
| **HME6** | W3110 *galK*$_{tyr145UAG}$ Δ*lacU169* [λ c*I857* Δ(*cro-bioA*)] | This strain contains the *galK* assay system for oligo recombination. |
| **HME63** | HME6 *mutS<>amp* | Defective for MMR thus gives high frequency oligo recombination. |
| **MSB1** | HME63 *araBAD<>Kan* | Lacks promoter for the *ara* genes and grows red when cultured on TA pla |
| **MSB2** | HME63 *dnaB:R74A* | Contains a *dnaB* point mutation |
| **MSB3** | HME63 *dnaB:R164A* | Contains a *dnaB* point mutation |
| **MSB4** | HME63 *dnaB:K180A* | Contains a *dnaB* point mutation |
| **MSB5** | HME63 *dnaB:R328/9A* | Contains a *dnaB* point mutation |
| **CM742** | *E. coli* K-12 d*naA46(ts)* | Contains a temperature sensitive *dnaA* mutation, CGSC# 12549 |

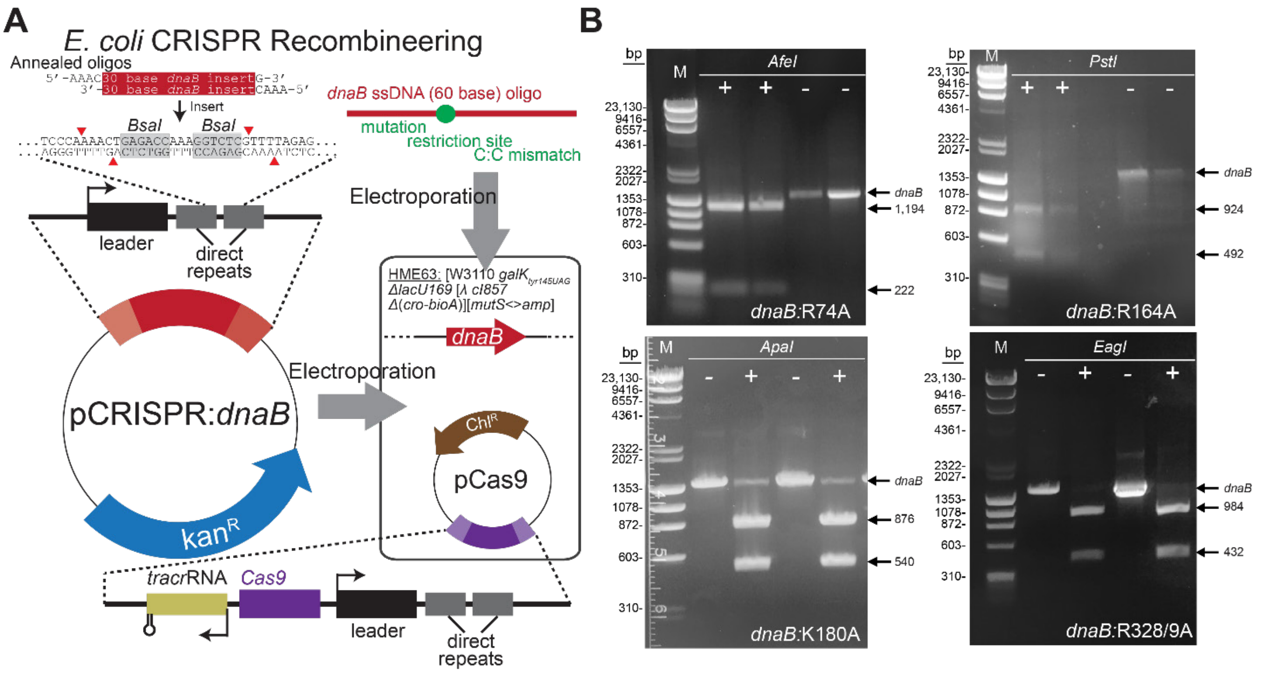


**Fig. S1 CRISPR-Cas9 recombineering to create *dnaB:muts***

(**A**) CRISPR-Cas9 recombineering using the dual-plasmid system. A target gRNA for the desired mutation site on the *dnaB* gene was inserted into the *BsaI* cloning sites of pCRISPR, before electroporating both the pCRISPR plasmid and the recombination DNA oligonucleotide, engineered to contain the mutation, a novel restriction enzyme site for screening, and a point mutation to disrupt the PAM (5'-NGG) sequence. (**B**) Restriction digest gels for each of the engineered *dnaB* mutants showing successful digest at the novel restriction site and confirming *dnaB* gene mutation.

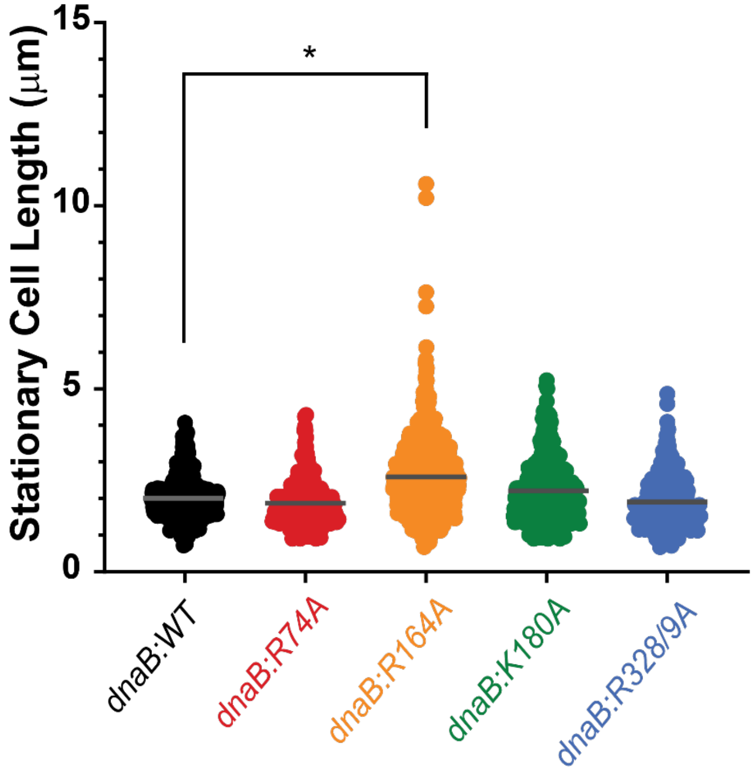


**S2 Fig. Quantification of stationary phase *dnaB:mut* cell length**

Overnight stationary phase cultures were imaged by microscopy, and cell length was measured by blinded visual quantification. Average cell length is represented by the black bar in the middle of the data set. Average length in order from left to right: 2.0 ± 0.5 μm, 1.9 ± 0.6 μm, 2.6 ± 0.9 μm, 2.2 ± 0.8 μm, and 1.9 ± 0.7 μm. n ≥ 400 events. Black bars above graph indicate statistically significant differences, where p-values are *< 0.05.

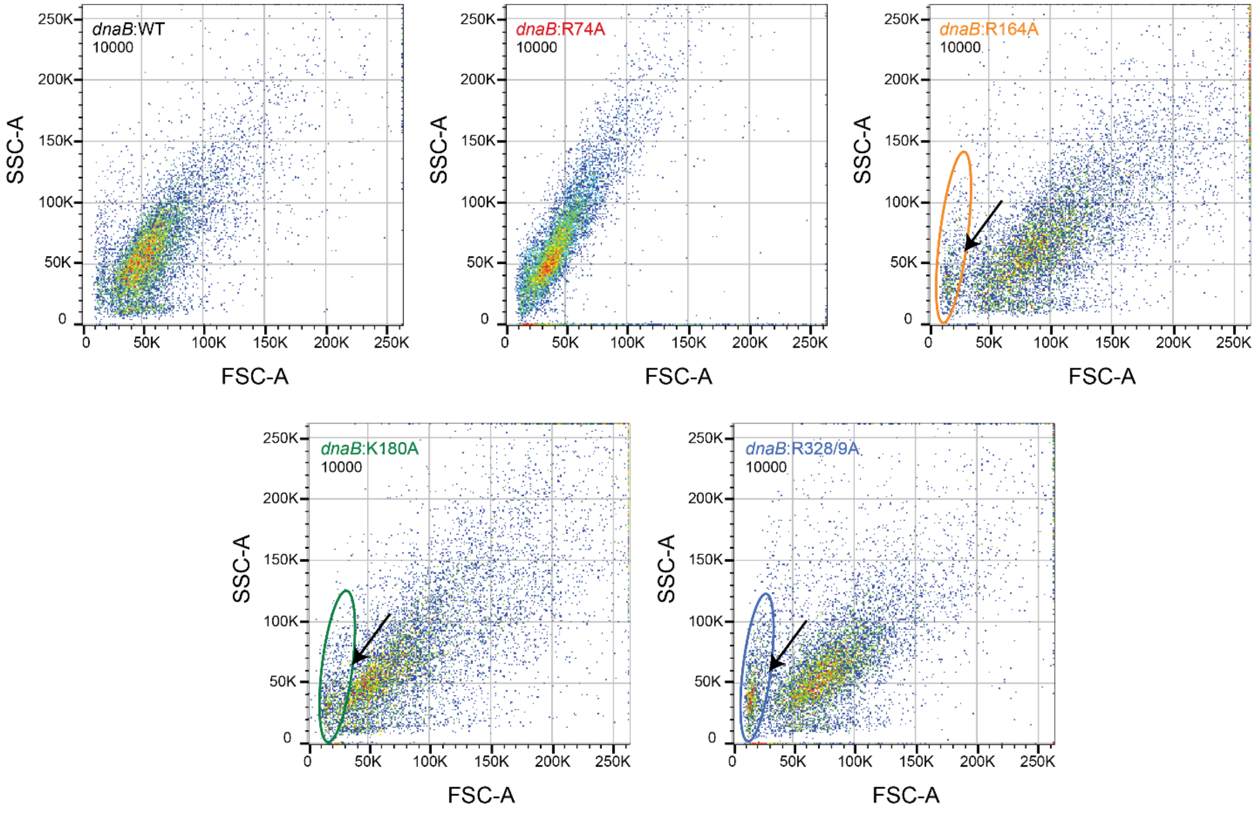


**S3 Fig. *dnaB:mut* strains show changes in cell size and complexity populations**

Cell cultures exposed to rifampicin and cephalexin were then analyzed by FACS. The forward (FSC) and side scatter (SSC) gains were set based on the parental strain, and then scatter plot data was collected for all strains, n = 10,000 events. Dense populations of cells are indicated with red, moderate with green, and minor or diffuse population with blue. The parental strain has a single cell population that is primarily green. *dnaB:R74A* has a single population that is the most concentrated of all the strains (including parental), signified by strong red signal. *dnaB:R164A* is significantly more diffuse than the parental strain and has a small secondary population (indicated by circle/arrow ). *dnaB:K180A* and *dnaB:R328/9A* both have primary populations similar to that of the parent but also show a small secondary population (circle/arrow).

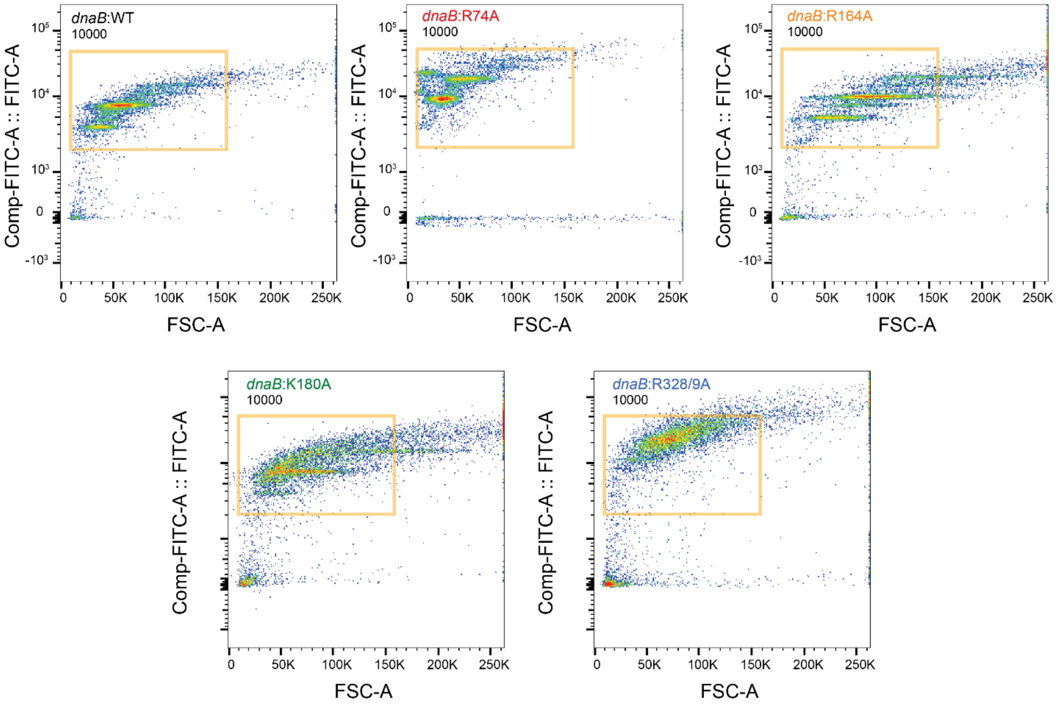


**S4 Fig. Rifampicin 'run-out' FACS reveals unique concentrations of DNA per cell size for *dnaB:mut* strains.**

FITC vs FSC scatter plot of cell cultures exposed to rifampicin and cephalexin analyzed by FACS. FITC indicates the chromatin staining intensity, and FSC indicates cell size. Dense populations of cells are indicated with red, moderate with green, and minor or diffuse population with blue. Fixed yellow box is intended to highlight shift in the location of populations. The parental strain has two major cell populations, with cell size increasing with chromatin. *dnaB:R74A* is shifted to the left, with small cells containing large amounts of chromatin. *dnaB:R164A* has three elongated FITC populations, meaning that a single concentration of chromatin is contained within a wide range of cell sizes. The cell populations of *dnaB:K180A* have lost definition, diffusing into one another. *dnaB:R328/9A* only has a single large population near the top of the FITC axis, indicating that this strain contains almost exclusively varied cell sizes with large amounts of chromatin.

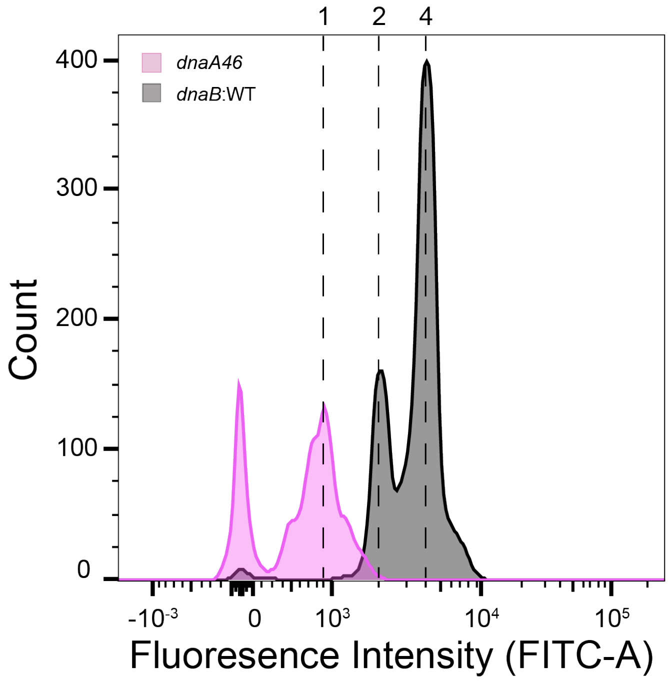


**S5 Fig. Actively growing parental (*dnaB:WT*) cells have major peaks for 2 and 4 chromosomes**

A control experiment measuring chromosome density for the parental strain *dnaB:WT* and a single-chromosome strain, *dnaA46(ts)*. Chromosome density was measured by flow cytometry (FACS) in log phase rifampicin ‘run-out’ cultures stained with Sytox green (n = 10,000 events). Chromosome integers are indicated at the top of the graph.

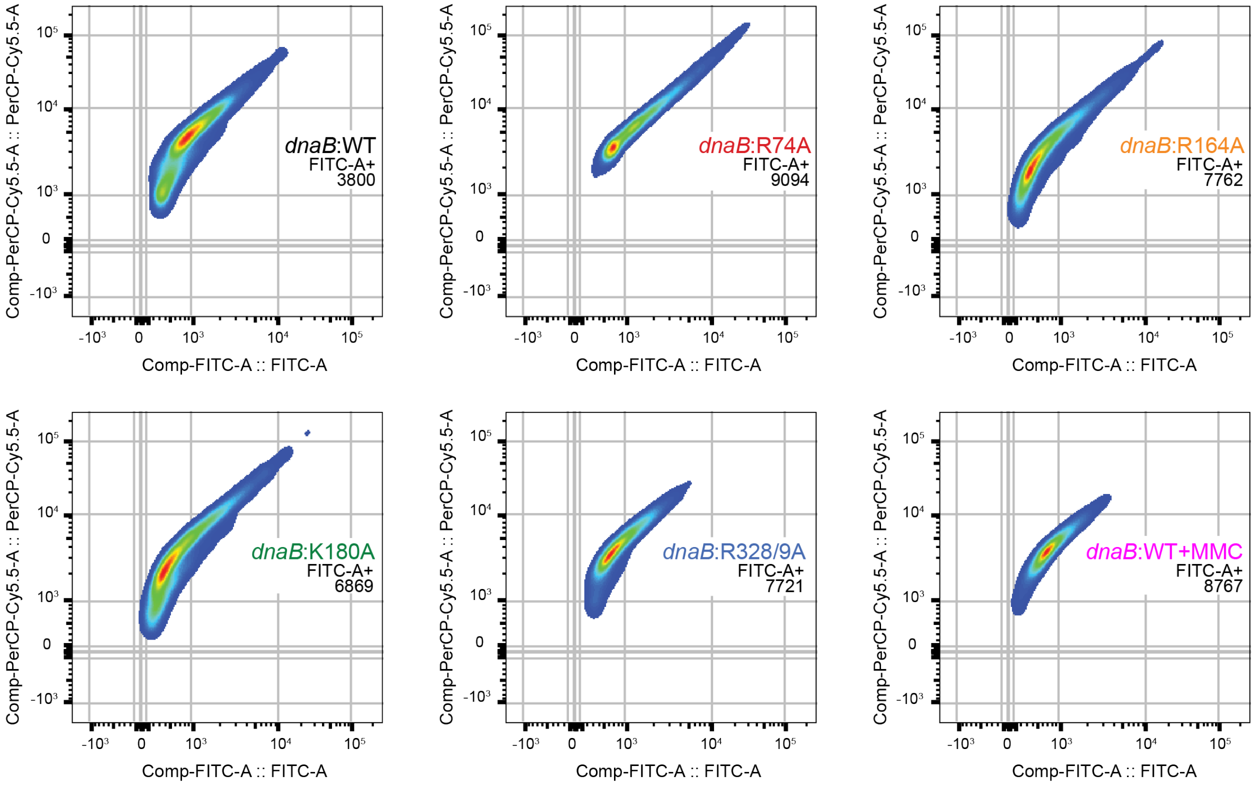


**S6 Fig. FACS data for *dnaB:mut* TUNEL assay**

Flow cytometry data of log phase cells plotted to show the relationship between DNA breaks and total amount of DNA. In these smoothed density plots, red represents concentrated cell populations, and dark blue represents highly diffuse cell populations. The grid is present to highlight changes in the size or location of cell populations. Small overall areas indicate concentrated populations that have a uniform and consistent distribution of DNA breaks (as that seen for parent + MMC). Noticeable tailing relative to the parental strain represents populations of cells that have increased DNA and DNA damage (as seen for *dnaB:R74A*). Y-axis shifts indicate changes in DNA repair or damage sensitivity; while X-axis shifts indicate changes in the amount of chromatin.

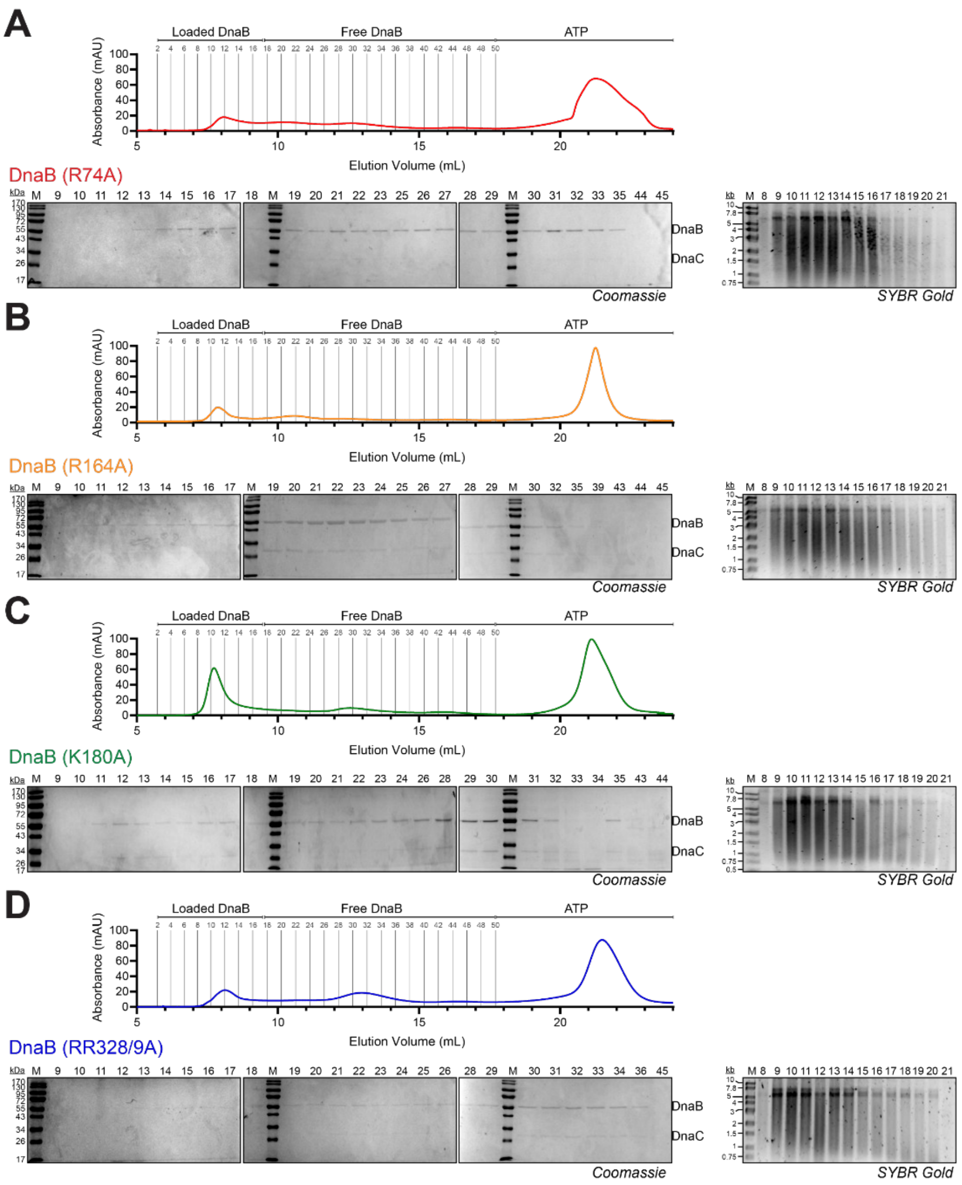


**S7 Fig. Size exclusion chromatography loading assay for the DnaB mutants**

DnaB **(A)** R74A, **(B)** R164A, **(C)** K180A, and **(D)** R328/9A were preincubated with DnaC, M13, and ATP before injecting onto a preequilibrated S200 10/30 size exclusion column according to the Materials and Methods. Example chromatogram and associated SDS-PAGE (Coomassie) and agarose (SYBR-Gold) gels used to monitor loaded DnaB and Free DnaB areas.

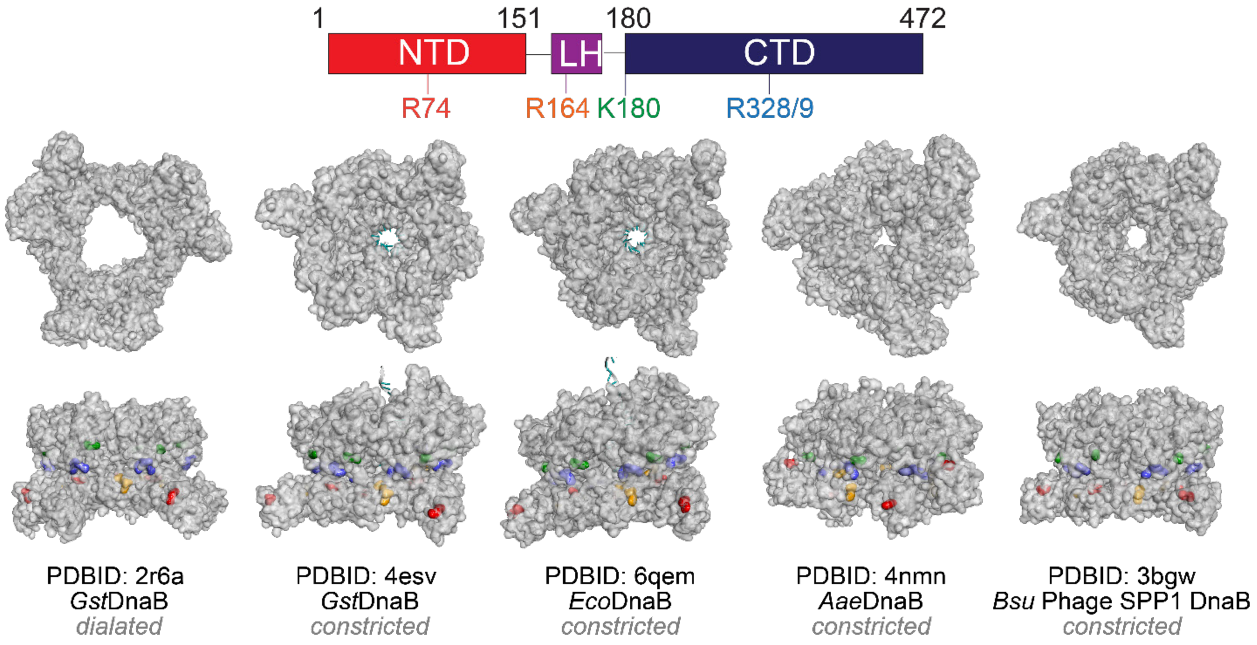


**S8 Fig. Constricted Crystal of *E. coli* DnaB with mutants mapped.**

Crystal structure of dilated DnaB (PDB: 2r6a), constricted cracked DnaB (lockwasher, PDB: 6qem), and constricted DnaB (PDB: 3bgw) with mutated residues highlighted: R74A in red, R164A in orange, K180A in green, and R328/9A in blue. The linear protein map (top) shows the location of each mutation relative to functional domains.
